# OmniExtract: An automatic data extraction tool based on Large Language Model and Prompt Engineering

**DOI:** 10.1101/2025.09.11.675332

**Authors:** Yibo Wang, Bixia Tang, Sicheng Wu, Yuyan Meng, Demian Kong, Wenming Zhao

**Affiliations:** National Genomics Data Center, China National Center for Bioinformation, Beijing 100049, China; Beijing Institute of Genomics, Chinese Academy of Sciences, Beijing 100049, China; University of Chinese Academy of Sciences, Beijing 100049, China

## Abstract

Extracting structured information from documents or scientific papers is crucial for data sharing and retrieval. Recently, Large Language Model (LLM) has shown its impressive ability in text understanding and several tools based on LLM has been developed. However, it’s still difficult to find a universal and user-friendly tool for various practical extraction tasks. To address this challenge, we propose OmniExtract, an automatic data extraction tool with user-friendly configuration files which can adapt to various data extraction tasks. OmniExtract uses a prompt optimized engineering to improve prompt and obtain high performance, and it can support a comprehensive data extraction including text and tables. Evaluation results show that OmniExtract obtains a high accuracy over 80% for 3 datasets. Furthermore, two additional data extraction applications using OmniExtract have been provided, achieving an accuracy of 92.21% and an average F1 score of 0.83 respectively. The data reliability performance shows that OmniExtract is a valuable tool for database updating.

## Introduction

With the advancement of scientific research, a large amount of research results is published as papers or documents, which form the foundation of building domain-specific knowledgebases such as EWAS for epigenome research[1], LncRNAWiki for lncRNA research[2], and BioKA for biomarker research[3]. However, most of them are constructed depend on manual curation, making the knowledge mining process expensive and inefficient, and facing the problem of continuous maintaining. Therefore, using efficient tools to improve the efficiency of data mining has become an urgent demand.

Several methods such as DECaF, MaterialsBERT[4] have been developed for automatically extracting simple task from documents such as NER (Named Entity Recognition). These methods rely on hundreds to thousands of high-quality annotated data and are difficult to generalize to other information extraction tasks, which limits their applicability. The large language model such as ChatGPT[5] shows impressive ability in text understanding, and recently several LLM-based extraction methods have been developed such as AIE[6], SBERT[7], ChatExtract[8], HybridRAG[9], LLM-IE[10], MedPromptExtract[11], RUIE[12]. These methods can be broadly classified into three types including RAG (Retrieval-augmented Generation, RAG), Prompt Engineering, and fine-tuned model[13]. Compared with the method based on RAG and fine-tuned model, prompt engineering does not rely on a large amount of high-quality annotated data and can switch models used for information extraction, making it flexibly applicable to various information extraction tasks in different scenarios. However, some limitations still exist. Firstly, most of these methods are focus on specific extracted task and the code or implementation of most methods cannot be obtained, which is difficult transfer to other tasks. Secondly, open-source methods such as LLM-IE are dedicated to providing a package for building a module that extracts specific information from a given text, and cannot be directly applied to extract target information from raw documents. Furthermore, these methods mainly focus on extract text information and have limitation to handle extracting tables from supplementary materials. Therefore, there is still a gap in reality application for automatic information extraction which capable of extract multiple data types and can be simply transfer with user-friendly configuration.

Here, we present OmniExtract, a universal and automatic tool used for data extraction. OmniExtract is developed based on prompt engineering and large language model which has these advanced features: 1) It provides a user-friendly way to extract target information from the original documents by simply modifying configuration files; 2) It enables to use existing curation data to modify prompt and get an optimized prompt to extract information, which make it can adapt to various curation requirement; 3) It can extract information from text and table, which make it can fulfill most of the reality application requirements. OmniExtract is evaluated on three standard corpuses using 8 various LLM models, the results show OmniExtract achieve 81.75%-89.00% accuracy in 3 datasets. These features as well as the performance will make OmniExtract as a valuable tool for data extraction which can help to improve knowledge update efficiency.

## Results

### Overall Performance Evaluation

We first tested the extraction performance of OmniExtract using different models on the WikiReading Recycled dataset[14] and the Yidu-S4K (https://tianchi.aliyun.com/dataset/144419) dataset. After prompt optimization, the models achieved extraction accuracy rates of 77.9%-84.3% on the WikiReading Recycled dataset and 67%-89% on the Yidu-S4K dataset (Table 1). Most models with a scale exceeding 14B achieved extraction accuracy exceeding 80%, demonstrating the reliability of the extraction tool (considering the subjective nature inherent to information extraction tasks).

**Table 1.** Performance Evaluation Using WikiReading Recycled and Yidu-s4k dataset.

|  | WikiReading Recycled<br>Accuracy(improvement) | Yidu-s4k<br>Accuracy(improvement) |
| --- | --- | --- |
| Qwen2.5-7B-Instruct-1M | 79.10% (5.70%↑) | 70.33% (**38.00%**↑) |
| Meta-Llama-3.1-8B-Instruct | 80.30% (6.00%↑) | 67.00% (7.00%↑) |
| Qwen2.5-14B-Instruct-1M | 80.20% (5.40%↑) | 83.33% (12.00%↑) |
| Mistral-Small-24B-Instruct-2501 | 82.95% (3.25%↑) | 86.67% (3.67%↑) |
| Qwen2.5-32B-Instruct | 82.40% (2.20%↑) | 82.67% (0.33%↓) |
| Llama-3.3-70B-Instruct | 77.90% (0.00%) | 82.33% (5.33%↑) |
| Qwen2.5-72B-Instruct | 82.50% (0.30%↑) | **89.00%** (6.33%↑) |
| Mistral-Large-Instruct-2407 (130B) | **84.30%** (**6.90%**↑) | 83.00% (1.67%↑) |

↑ indicates the accuracy improvement achieved by using the optimized prompt compared to the unoptimized prompt; ↓ indicates the accuracy reduction after using the optimized prompt.

With the exception of the two models below 10B (Qwen2.5-7B-Instruct-1M and Meta-Llama-3.1-8B-Instruct), the extraction performance of the other models was generally similar. However, the highest accuracy rates on both tasks were achieved by models exceeding 70B (Mistral-Large-Instruct-2407 for the WikiReading Recycled dataset and Qwen2.5-72B-Instruct for the Yidu-S4K dataset). Furthermore, the extraction performance of the four different Qwen models showed an increasing trend with parameter count, suggesting that larger-scale models can achieve higher extraction accuracy ceilings, which is consistent with the understanding that model capabilities exhibit logarithmic growth with increasing parameters.

We further analyzed the magnitude of accuracy improvement achieved by the models after prompt optimization. Three models exceeding 70B showed an average improvement of 2.4% on the WikiReading Recycled dataset and 4.4% on the Yidu-S4K dataset. In contrast, 4 models below 24B exhibited larger gains, with average improvements of 5.7% on the WikiReading Recycled dataset and 19% on the Yidu-S4K dataset. Nonetheless, even for models larger than 70B, prompt optimization could still yield improvements exceeding 6 percentage points, which validates the efficacy of the prompt optimization module.

Next, we used the CriticalCoolingRates dataset[8] to test the models’ effectiveness in extracting multiple materials and their corresponding critical cooling rate values (with different units) from raw literature (Table 2). Most models performed poorly, achieving extraction precision below 60%. However, the Qwen2.5-72B-Instruct model achieved 82.63% precision and 71.88% recall. Considering the challenging nature of this task (original reports indicate both precision and recall were below 80%[8] – though not comparable, this demonstrates the task’s difficulty), these results demonstrate that our tool can achieve acceptable multi-entity extraction performance.

**Table 2.** Performance Evaluation Using CriticalCoolingRates dataset.

|  | **Precision** | **Recall** |
| --- | --- | --- |
| Qwen2.5-7B-Instruct-1M | 23.16% | 47.40% |
| Meta-Llama-3.1-8B-Instruct | 11.44% | 48.44% |
| Qwen2.5-14B-Instruct-1M | 55.31% | 66.84% |
| Mistral-Small-24B-Instruct-2501* | - | - |
| Qwen2.5-32B-Instruct | 46.74% | 70.83% |
| Llama-3.3-70B-Instruct | 44.13% | 64.58% |
| Qwen2.5-72B-Instruct | **82.63%** | **71.88%** |
| Mistral-Large-Instruct-2407 (130B) | 46.52% | 55.73% |

Mistral-Small-24B-Instruct-2501 does not have enough context length to handle the whole literature content.

### Application for the automatic extraction of multiple traits in dog breed standardization

Dogs are among the earliest domesticated animals and exhibit diverse phenotypes. Currently, there are more than 400 dog breeds worldwide [15], and 283 breeds of them have been registered in the AKC (American Kennel Club) with detailed standards for morphologic traits such as body weight, height, tail length, etc. These morphological traits have been selected through long-term artificial breeding, showing significant differences among different breeds and having a strong correlation with the genetic background of specific breeds. Therefore, these traits are valuable data resources for canine genetics research. To investigate the genetics of complex traits for dogs, GWAS

(Genome-wide association study) is used to research the association with genotype and trait using standard trait value, and several candidate genes have been found [16]. However, since the traits have been provided in the breed registered document, it will be inconvenient to retrieve the value of multiple traits for each breed. Thus, our goal is to automatically extract phenotypic information from breed standard documents and standardize some important phenotypes, in order to construct a phenotypic dataset convenient for canine GWAS research. OmniExtract is used to automatically extract traits from dog breed standardization files, and the whole extracted traits can be downloaded from iDog 2.0[17].

To optimize the prompt for extracting traits, a comprehensive dataset containing 39 distinct traits across 62 breeds was manually curated. A simple initial prompt for extracting traits has been constructed, after the prompt engineering tuning, an optimized prompt has been obtained as Figure 1.The used large language model Qwen2.5-72B-Instruct was configured, OmniExtract was applied to the breed standard files’ extraction. 100 breed standard files were selected for evaluating extraction accuracy. A total of 3,900 trait values were manually checked, resulting in an extraction accuracy rate of 89.62%.

**Figure 1.**
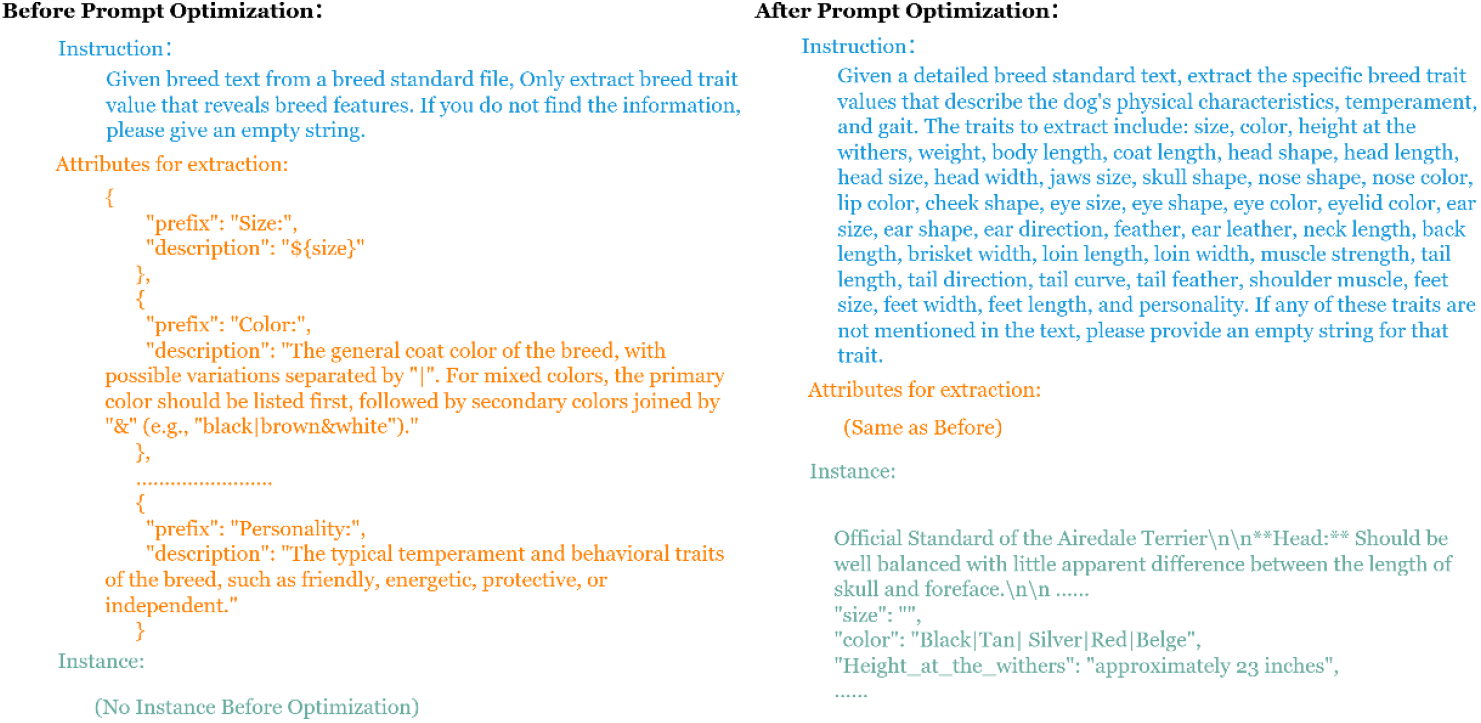
The initial prompt and optimized prompt used to extract trait values from breed standardization files.

Further, we analyzed the causes of extraction errors. The top five traits with the highest error rates were Head Width, Head Length, Size, Cheek Shape, and Weight (Figure 2). The reasons for errors in these five traits are respectively:

**Figure 2.**
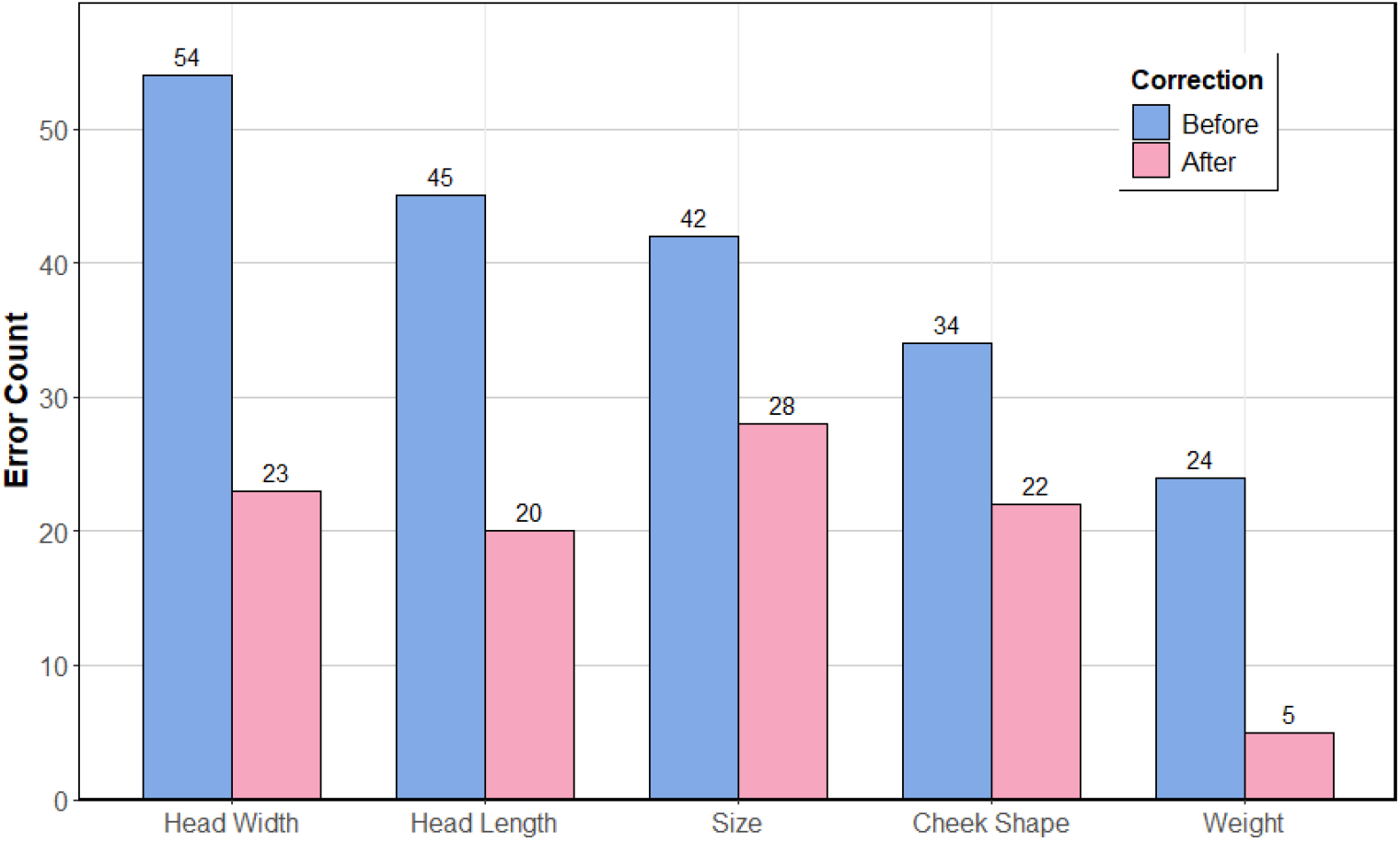
The top five traits with the highest error rates.

1. Semantic Confusion: For Head Width and Head Length, the erroneous extraction results often describe the length and width of the skull; for Cheek Shape, the erroneous results often describe the Muzzle (meaning the animal’s snout or nose). We observed that in certain breed standard documents (e.g., Golden Retriever), descriptions labeled as “Head” actually pertain to skull characteristics. Similarly, some documents’ “Skull” sections substantively describe head features. Furthermore, within breed standards, Head and Skull, as well as Cheek and Muzzle, frequently appear in close proximity within the same paragraph. This semantic and positional ambiguity likely contributes to the occurrence of such extraction errors.
2. Failure to meet formatting requirements: For Body Size, we expect outputs to be descriptive keywords such as small, medium, large, or combinations thereof. However, as these keywords are absent in certain breed documents, the model often extracts descriptions of height or weight in an effort to capture comprehensive information. This violates our formatting rules.
3. Generation without consulting the document: Many documents do not contain Weight information for specific dog breeds. However, the model may fabricate weight data based on knowledge from unknown sources. Some of these fabricated weights coincidentally match data from authoritative sources like AKC.

To verify the above analysis and further improve the extraction accuracy, we rewrote and optimized the prompts for these 5 traits and extracted the values of them. Compared with the previous results, the error rate of this extraction has decreased by 33.33%-79.17% (Figure 2). Combining the results of the two extraction processes, the extraction accuracy has risen to 92.21%

These results indicate that the initial prompt for the information extraction task requires more refined curating standard descriptions. This need arises because data-driven optimization processes struggle to align with implicit human intent – a likely reason why it is difficult to improve the accuracy rate to above 90% or 95% when using simple initial prompts.

### Application for comprehensive information automatic extraction in EWAS

Epigenome-Wide Association Study (EWAS) has become a powerful approach to identify epigenetic variations associated with biological traits[18]. To provide a comprehensive collection of high quality EWAS associations in support of systematic investigations of complex molecular mechanisms associated with different biological traits, a manually curated knowledgebase EWAS atlas has been constructed and published in 2018[1]. For more than 6 years, EWAS atlas has consistently maintain and update the database by manually curated information which is time-consuming and hard-working. To improve the data update efficiency, OmniExtract was used to automatic extract comprehensive information including trait, cohort, and EWAS association information which involves text and table extraction from main text and supplemental materials.

We first apply the OmniExtract tool to the task of extracting trait names and trait types from the abstracts of EWAS-related literature. An EWAS study may involve multiple traits, and according to the requirements of the EWAS Atlas database, these traits need to be classified into five types: cancer, non-cancer disease, phenotype, environmental factor, and behavior. In addition, some traits that are not the target of EWAS research need to be accurately filtered out, which further increases the difficulty of information extraction. We utilized the optimized prompt to assist the extraction task and manually checked the extraction results for 100 articles. This process yielded a precision of 88.36%, a recall of 87.76%, and an F1 score of 0.8705. The majority of extraction errors stem from either incorrectly extracting non-target traits or failing to extract target traits, as exemplified in Figure 3.

**Figure 3.**
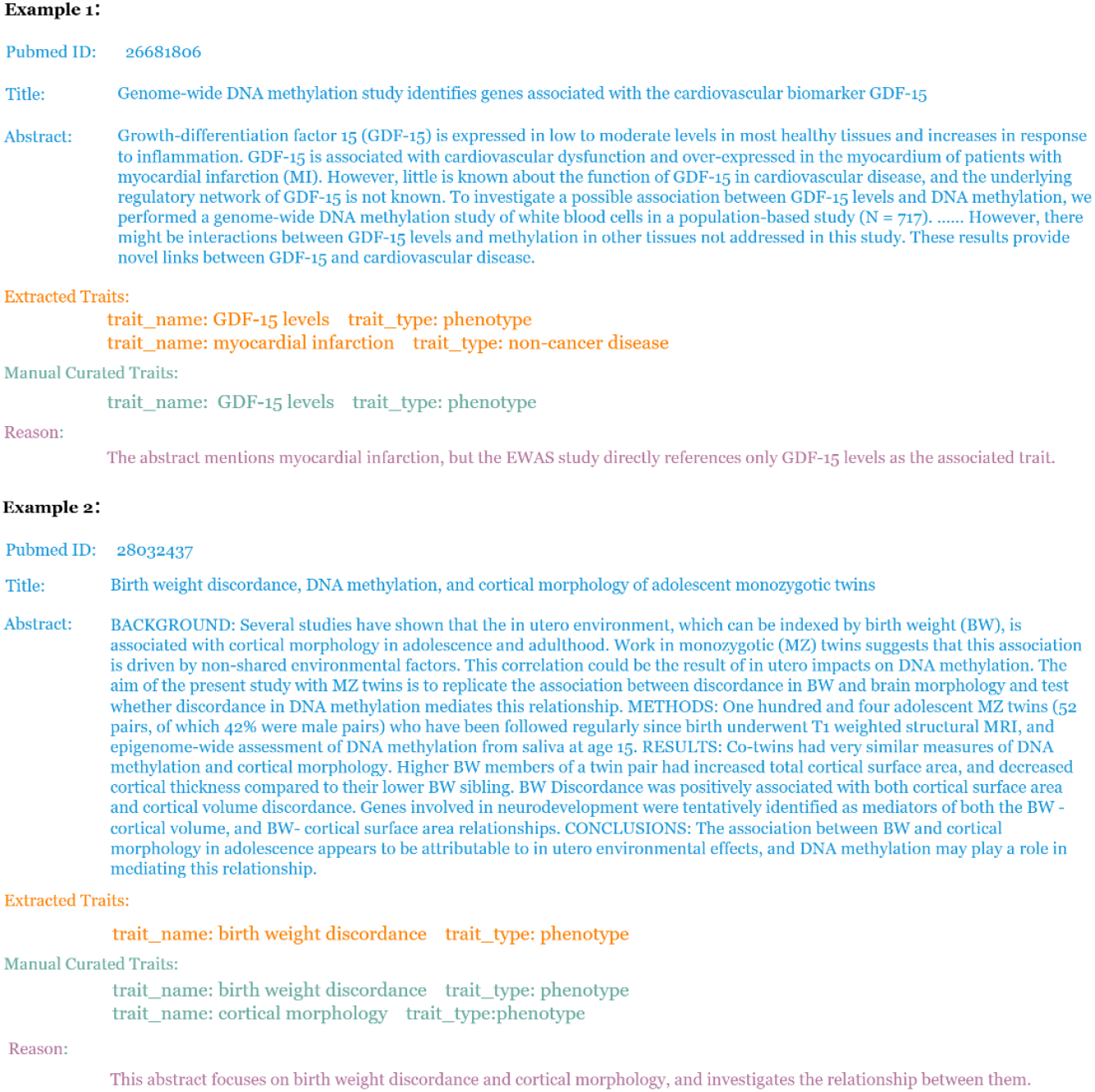
Error cases for the target traits extraction.

Further, we utilized the OmniExtract tool to extract cohort metadata (sample size, age range, male proportion, platform, cohort name, ancestry, etc.) from EWAS full-text literature. Considering the complexity of population cohort information and the variable number of cohorts that may be involved in a single EWAS study, this constitutes an exceptionally challenging task. After manual verification, the OmniExtract tool ultimately achieved a precision of 81.75%, recall of 74.68%, and an F1 score of 0.7806, which are comparable to previous test outcomes on the CriticalCoolingRate dataset. We analyzed the causes of the lower recall rate and identified that two articles contained a significantly higher-than-average number of cohorts (16 and 17 cohorts for these two articles, with 5.03 and 5.43 z-scores). These cohorts were erroneously omitted during extraction, accounting for 36.7% of all missed cohorts (Figure 4). Such corner cases constitute the primary factor for recall rate degradation in similar tasks, while the decline in precision is predominantly attributable to factual errors stemming from the high complexity of information extraction (Figure 5).

**Figure 4.**
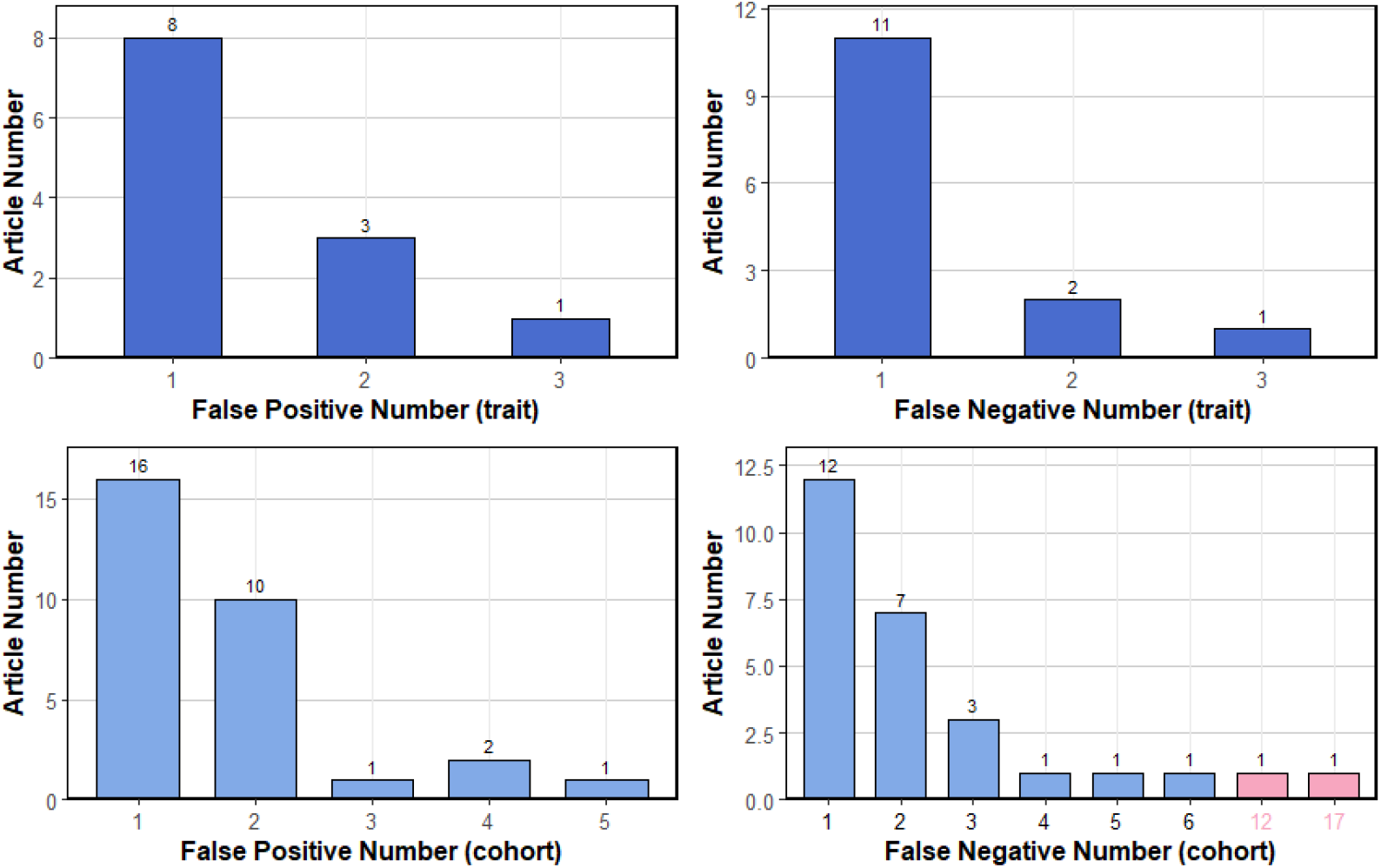
Distribution of article numbers about error counts of the trait and cohort extraction results

**Figure 5.**
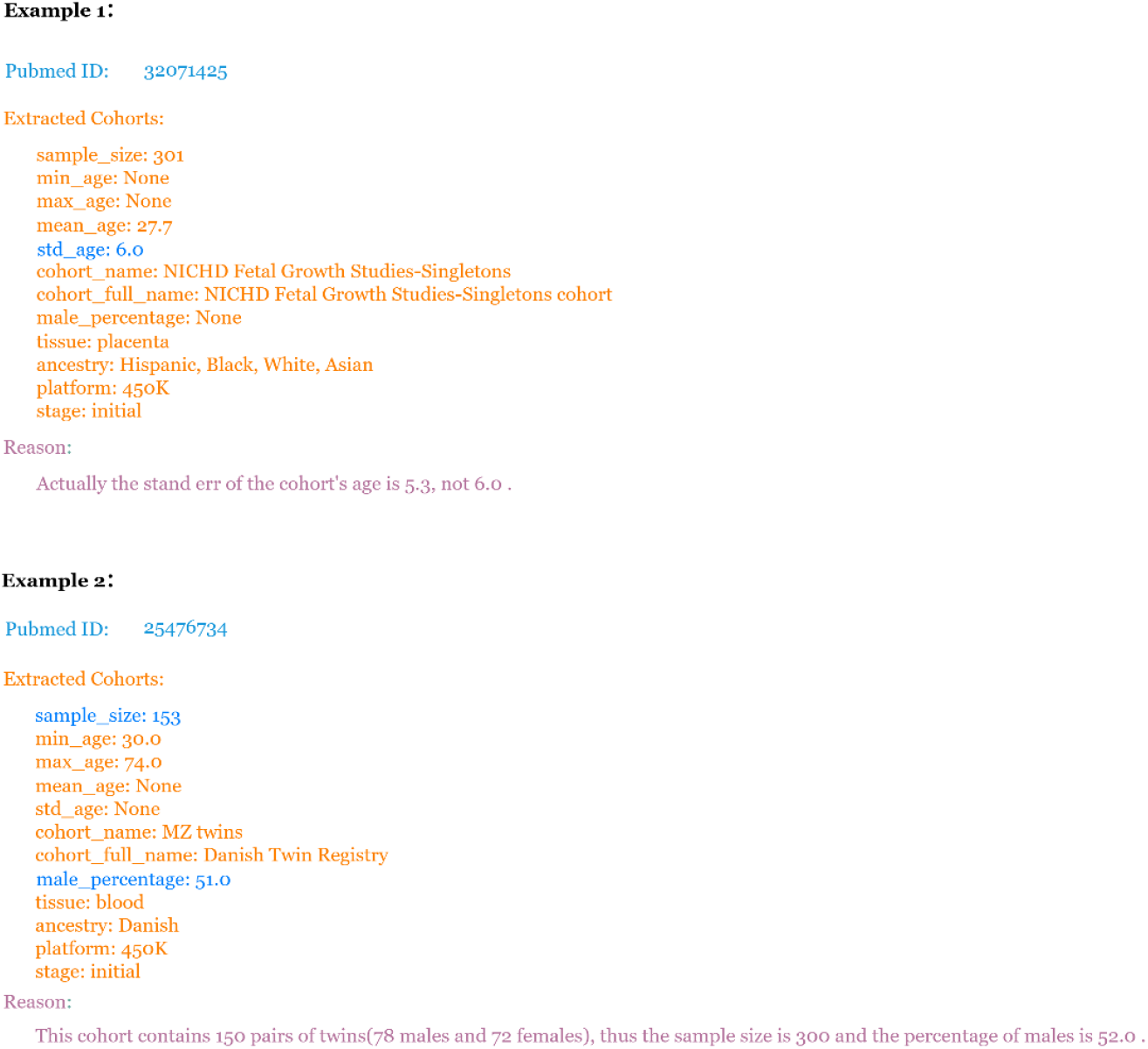
Error cases of mistakenly extracted cohort.

The extraction of EWAS association data was completed using OmniExtract’s table processing module. We processed 39 tabular files (xlsx/xls/csv/tsv) containing 116 tables from supplementary materials across 21 publications. Manual verification confirmed that 86.21% of the 116 tables could be classified and extracted correctly. The primary cause of errors stems from the limited information provided by table field names, making it challenging to map them to the intended extraction targets in certain scenarios.

## Discussion

OmniExtract provides a one-step solution for extracting multi-property entities from original documents and tables, encompassing all the functions of file parsing, prompt optimization, and information extraction. OmniExtract has two features: (1) User-friendly: Users can utilize the corresponding functions by modifying predefined YAML file templates. (2) Stable performance: Using mainstream closed-source or open-source models, the extraction accuracy of multi-property entity extraction tasks can reach more than 80%, and can reach more than 90% after targeted adjustments. In addition to switching to more capable models, the extraction performance of the tool can still be enhanced by better aligning with users’ implicit criteria. The following are some ideas for improving extraction performance, which will be implemented in future versions:

1. Collaborating with LLM as a corrector: As demonstrated in the extraction of cohort information from EWAS literature, the main factor contributing to decreased recall is information loss when handling atypical tasks, while the reason for decreased precision is some factual errors. Therefore, after information extraction, we can use capable models to complete verification and correction for these two situations. Adopting this strategy of small model extraction followed by large model correction can ensure extraction efficiency while improving extraction accuracy.
2. Implementation of Context Compression Technology: In the current version, text extracted from raw documents is just roughly chunked before being directly fed into the model for extraction. This approach not only incurs higher computational costs but also allows excessive context length to adversely affect extraction effectiveness. Therefore, in future releases, we plan to integrate context compression technology into the workflow to minimize the impact of irrelevant information on model extraction. Context compression technology streamlines the content input to models by filtering out irrelevant documents or summarizing the documents, which can improve the reliability of answers costs while reducing computational costs. We plan to introduce this technology in the future, using an independent small model to filter out content irrelevant to the extraction target before performing information extraction, thereby enhancing the effectiveness of information extraction.

Additionally, our table extraction feature currently focuses primarily on extracting information from table files, and has not optimized the workflow for extracting table information from PDF files. In future versions, we plan to integrate parsing tools with stronger PDF table parsing capabilities and optimize the table extraction process from PDF files to achieve better extraction results.

## Methods

### System design

Our goal is to build an information extraction tool that is (1) functionally complete: this tool needs to be able to conveniently extract information from original files without introducing additional tools or modules, (2) user-friendly: users only need to modify the configuration file to use the tool for information extraction, and (3) stable in performance: the tool can achieve good performance when completing different extraction tasks. To achieve the aforementioned goals, we have constructed the information extraction tool named OmniExtract using DSPy[19], which is a declarative, modular framework for building AI applications that offers efficient prompt optimization functionalities. OmniExtract seamlessly integrates Marker(https://github.com/datalab-to/marker), a document parsing tool, to provide robust support for processing raw PDF documents. OmniExtract comprises four modules: (1) Raw document processing, (2) prompt optimization based on existing data, (3) parallel information extraction & assessment, (4) table extraction workflow. We provide highly encapsulated APIs for each module, enabling the complete process from raw documents to information extraction through simple configuration file modifications.

**Figure 6.**
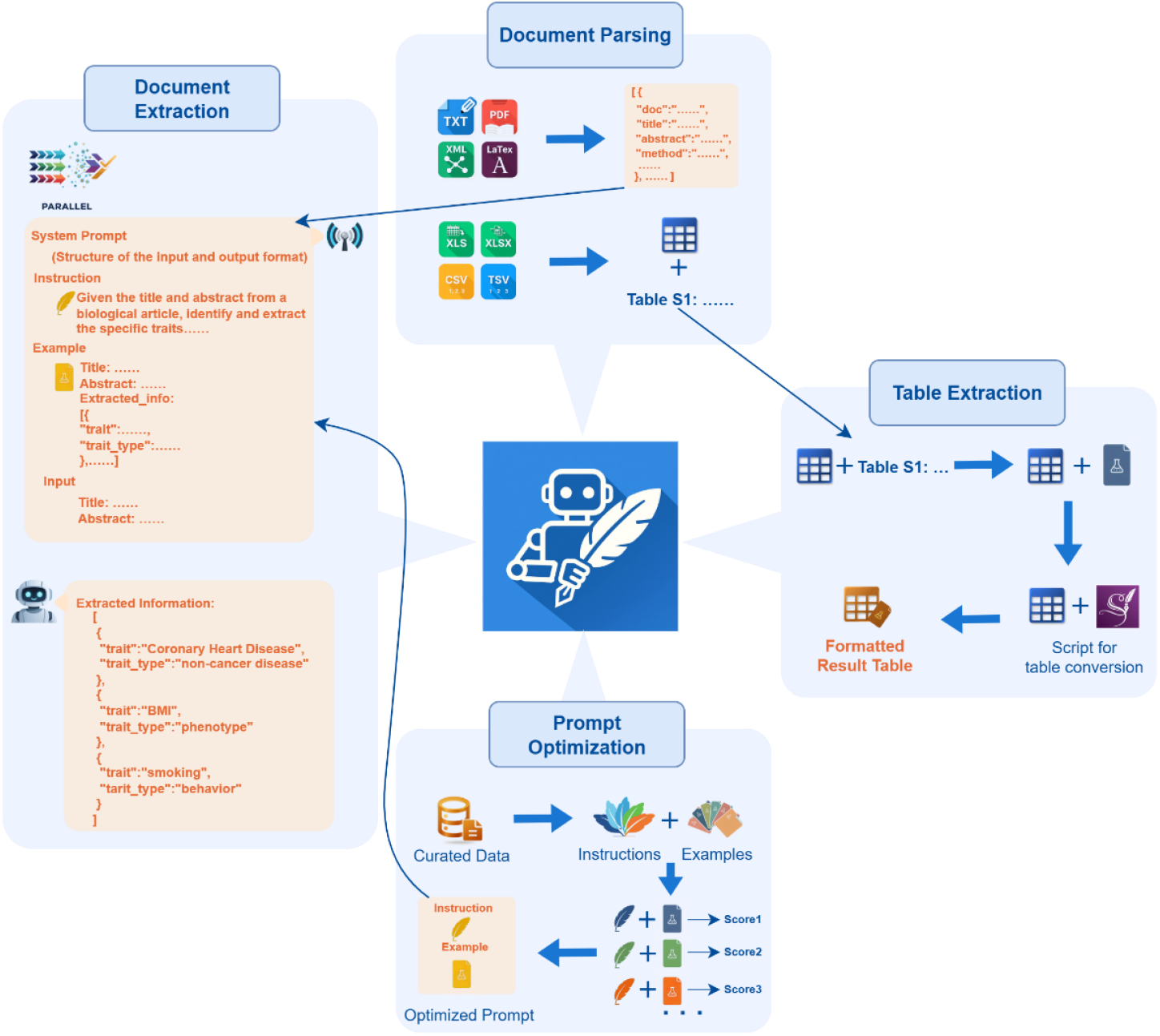
System design of OmniExtract.

### raw document processing

For general literature documents (typically in PDF format), we have integrated the Marker document parsing tool, which can quickly and accurately parse documents into LLM-friendly markdown format for subsequent information extraction. Additionally, we have developed a rule-based parsing pipeline for XML format files provided by PubMed Central (PMC) and ScienceDirect, as well as TeX format files from arXiv, which can parse the main content of documents into Markdown format. Furthermore, we have provided a text segmentation method. This method matches section headings within the literature using keyword matching, dividing the document into sections such as the Title, Abstract, Introduction, Methods, and Results. The segmented documents are saved in JSON format. This segmentation provides more precisely scoped context for subsequent information extraction.

For table information extraction, we have also implemented rule-based parsing pipelines. These pipelines clean and convert tabular files (xlsx, xls, csv, tsv) by removing blank rows and separating table titles. The table content is stored in TSV format, while the table titles are saved to a metadata file. Regarding literature sourced from PubMed Central (PMC) and ScienceDirect, we have also customized parsing methods to extract the table into a TSV file.

### prompt optimization based on existing data

To improve the performance of information extraction, we have also encapsulated the prompt optimization workflow. We have modified the prompt optimization method miproV2[20] offered by DSPy to better adapt to information extraction tasks. We have also provided both rule-based and predefined AI evaluation methods to assess the performance of selected prompts during prompt optimization. Based on the provided curated data, users can automatically complete the prompt optimization process by supplying a configuration similar to the information extraction workflow and selecting an appropriate evaluation method.

DSPy offers several prompt optimization methods. Among them, miproV2[20] can generate few-shot examples and candidate prompts, and select the optimal combination via Bayesian search. This allows it to improve the extraction accuracy by properly utilizing the curated information. However, information extraction tasks tend to directly extract information from given text without requiring a long processing chain. Models can often generate roughly correct answers without needing rejection sampling. Moreover, the extracted information often has implicit format and content requirements, which can be reflected in curated results. Therefore, we modified the miproV2 method and used the curated results instead of few-shot examples generated by LLM for the downstream prompt generation and composition optimization steps.

Prompt optimization methods require providing metrics to evaluate generated results, in order to select the highest-scoring prompts based on existing data. We provide two methods to evaluate the effectiveness of prompt-example combinations:

1. Rule-based Method: We establish a set of rules to evaluate the degree of match between extracted results and reference results. For string-type attributes: Building upon the evaluation methods provided by DSPy, we additionally employ the Smith-Waterman algorithm[21], Ratcliff-Obershelp algorithm[22], Jaccard index[23], and LCS algorithm, taking the highest score as the string similarity metric. For other non-list/non-string attributes: We require the extracted result to exactly match the reference answer. For list-type attributes: We calculate the Jaccard similarity as the score.
2. Predefined AI Evaluation Method: We use an LLM to score the degree of match between the extracted results and the reference results.

Users can utilize the OmniExtract tool to process curated data and the matched original documents, building datasets for prompt optimization. Based on the modified MiproV2 method, the instruction-instance combination could be optimized to enhance the effectiveness of information extraction.

### parallel information extraction & assessment

We have encapsulated a parallel information extraction workflow based on the declarative features of the DSPy framework. This approach transforms the coding effort for building the workflow into configuring input content and the names, types, and descriptions of target information to be extracted. Combined with predefined or optimized prompts, this generates modules required for information extraction. We leverage DSPy’s built-in ParallelExecutor class to process extraction tasks concurrently. Furthermore, adhering to the “LLM as a judge”[24] paradigm, we predefine a scoring component to evaluate extraction results. Users can employ a different model (or the same model used for extraction) to assign confidence scores to the results, enabling efficient filtering of low-quality outcomes.

### table extraction workflow

Extracting information from tables differs from extracting from ordinary text: tables are more structured and have higher information density compared to ordinary text, typically yielding more extracted information. Considering these characteristics, we have designed a dedicated workflow for table extraction. After parsing the table, we transform the original table into a structured table containing extracted information through four steps: (1) table classification: determine whether the table contains target information based on LLM, (2) example table generation: extract target information from the first 10 rows of the table, store it as an example file, (3) script generation for table transformation: generate the code to convert the table into the example table format based on a coder model, and (4) convert the table into the specified format containing target information: run the script generated in step 3 and check the result. Users need to specify the prompt words for table classification and example table generation in the configuration file to complete this workflow. Similarly, the extraction performance of this process can be enhanced through the prompt optimization module.

### Datasets and evaluation metrics

Three public datasets were used to evaluate the information extraction performance of the tool:

1. WikiReading Recycled dataset[14]: This is a dataset designed for multi-property extraction tasks built from the Wikipedia information. We extracted information for 200 professional athletes from this dataset (including 12 attributes such as name, gender, country, and date of birth) for evaluation. This dataset is used to evaluate the performance of extracting multiple attributes of one entity from a short document.
2. Yidu-S4K dataset (https://tianchi.aliyun.com/dataset/144419): This is a Chinese electronic medical record dataset that includes tasks for medical entity and attribute extraction. We extracted 200 records from the training set of this task for evaluation. This dataset is used to evaluate the effectiveness of multi-property extraction for Chinese.
3. the CriticalCoolingRates dataset[8]: we randomly sampled records from 200 articles from the “MANUAL_raw” section, which containing materials’ names and their corresponding critical cooling rates. This dataset is used to evaluate the performance of extracting multiple entities from original documents.

For the first two datasets, we extract multiple attributes of individual entities from each piece of data and calculate the accuracy rate of the extracted attributes. For the third dataset, we adopt a “two-step” strategy: first parsing the PDF documents of the literature and extracting key paragraphs containing the material’s critical cooling rate, then combining the full text with these key paragraphs to extract the material names and critical cooling rates. We manually verified the extraction results by benchmarking against the dataset and calculated the precision and recall.

Furthermore, we demonstrate the application of our information extraction tool in two real-world scenarios:

1. Extracting multiple trait information from dog breed standard documents from AKC.
2. Extracting trait and cohort information from Epigenome-Wide Association Study (EWAS) literature, and capturing EWAS association data from table files.

### Models used for evaluation

We selected 8 open-source models from three mainstream model series (Llama, Qwen, Mistral) to evaluate tool information extraction capabilities across different model scales: Meta-Llama-3.1-8B-Instruct[25], Llama-3.3-70B-Instruct[25], Qwen2.5-7B-Instruct-1M[26], Qwen2.5-14B-Instruct-1M[26], Qwen2.5-32B-Instruct[27], Qwen2.5-72B-Instruct[27], Mistral-Small-24B-Instruct-2501(https://huggingface.co/mistralai/Mistral-Small-24B-Instruct-2501), and Mistral-Large-Instruct-2407 (https://huggingface.co/mistralai/Mistral-Large-Instruct-2407). Using vllm[28] (version 0.6.6), we deployed the models as a standalone service on a 4×A100 machine to provide interfaces for the extraction tool usage.

For the two application cases we provided, we used different models for processing: Qwen2.5-72B-Instruct was applied to trait information extraction of dog breeds. When extracting EWAS-related information, considering the complexity of the task, qwen-max was used for information extraction.

## Code Availability

The source code for OmniExtract is available on Github https://github.com/wyb39/OmniExtract.

## References

1. Li, M., et al., EWAS Atlas: a curated knowledgebase of epigenome-wide association studies. Nucleic Acids Res, 2019. 47(D1): p. D983–D988.

2. Liu, L., et al., LncRNAWiki 2.0: a knowledgebase of human long non-coding RNAs with enhanced curation model and database system. Nucleic Acids Res, 2022. 50(D1): p. D190–D195.

3. Wang, Y., et al., BioKA: a curated and integrated biomarker knowledgebase for animals. Nucleic Acids Res, 2024. 52(D1): p. D1121–D1130.

4. Yoshitake, M., et al., MaterialBERT for natural language processing of materials science texts. Science and Technology of Advanced Materials: Methods, 2022. 2(1): p. 372–380.

5. Welsby, P. and B.M.Y. Cheung, ChatGPT. Postgrad Med J, 2023. 99(1176): p. 1047–1048.

6. Yue, C., et al., Extract Information from Hybrid Long Documents Leveraging LLMs: A Framework and Dataset. arXiv, 2024.

7. Liu, Y. and S. Li, AutoIE: An Automated Framework for Information Extraction from Scientific Literature. arXiv, 2024.

8. Polak, M.P. and D. Morgan, Extracting accurate materials data from research papers with conversational language models and prompt engineering. Nat Commun, 2024. 15(1): p. 1569.

9. Sarmah, B., B. Hall, and R. Rao, HybridRAG: Integrating Knowledge Graphs and Vector Retrieval Augmented Generation for Efficient Information Extraction. arXiv, 2024.

10. Hsu, E. and K. Roberts, LLM-IE: A Python Package for Generative Information Extraction with Large Language Models. 2024.

11. Srivastava, R., et al., MedPromptExtract (Medical Data Extraction Tool): Anonymization and Hi-fidelity Automated data extraction using NLP and prompt engineering. arXiv, 2024.

12. Liao, X., et al., RUIE: Retrieval-based Unified Information Extraction using Large Language Model. arXiv, 2025.

13. Garcia, G.L., et al., A Review on Scientific Knowledge Extraction using Large Language Models in Biomedical Sciences. arXiv, 2024.

14. Dwojak, T., et al., From Dataset Recycling to Multi-Property Extraction and Beyond. 2020.

15. Ostrander, E.A., et al., Demographic history, selection and functional diversity of the canine genome. Nat Rev Genet, 2017. 18(12): p. 705–720.

16. Plassais, J., et al., Whole genome sequencing of canids reveals genomic regions under selection and variants influencing morphology. Nat Commun, 2019. 10(1): p. 1489.

17. Liu, Y., et al., iDog: a multi-omics resource for canids study. Nucleic Acids Res, 2025. 53(D1): p. D1039–D1046.

18. Rakyan, V.K., et al., Epigenome-wide association studies for common human diseases. Nat Rev Genet, 2011. 12(8): p. 529–41.

19. Khattab, O., et al. DSPy: Compiling Declarative Language Model Calls into Self-Improving Pipelines. 2023. 2310.03714 DOI: 10.48550/arXiv.2310.03714.

20. Opsahl-Ong, K., et al. Optimizing Instructions and Demonstrations for Multi-Stage Language Model Programs. 2024. 2406.11695 DOI: 10.48550/arXiv.2406.11695.

21. Smith, T.F. and M.S. Waterman, Identification of common molecular subsequences. J Mol Biol, 1981. 147(1): p. 195–7.

22. Ratcliff, J.W. and D. Metzener, Pattern matching: The gestalt approach. 1988.

23. Jaccard, P., Etude de la distribution florale dans une portion des Alpes et du Jura. Bulletin de la Societe Vaudoise des Sciences Naturelles, 1901. 37: p. 547–579.

24. Zheng, L., et al. Judging LLM-as-a-Judge with MT-Bench and Chatbot Arena. 2023. 2306.05685 DOI: 10.48550/arXiv.2306.05685.

25. Grattafiori, A., et al. The Llama 3 Herd of Models. 2024. 2407.21783 DOI: 10.48550/arXiv.2407.21783.

26. Yang, A., et al., Qwen2.5-1M Technical Report. 2025.

27. Qwen, et al. Qwen2.5 Technical Report. 2024. 2412.15115 DOI: 10.48550/arXiv.2412.15115.

28. Kwon, W., et al., Efficient Memory Management for Large Language Model Serving with PagedAttention. 2023.

